# Optimized iLID membrane anchors for local optogenetic protein recruitment

**DOI:** 10.1101/2020.09.05.268508

**Authors:** Dean E. Natwick, Sean R. Collins

## Abstract

Optogenetic protein dimerization systems are powerful tools to investigate the biochemical networks that cells use to make decisions and coordinate their activities. These tools, including the improved Light-Inducible Dimer (iLID) system, offer the ability to selectively recruit components to subcellular locations, such as micron-scale regions of the plasma membrane. In this way, the role of individual proteins within signaling networks can be examined with high spatiotemporal resolution. Currently, consistent recruitment is limited by heterogeneous optogenetic component expression, and spatial precision is diminished by protein diffusion, especially over long timescales. Here, we address these challenges within the iLID system with alternative membrane anchoring domains and fusion configurations. Using live cell imaging and mathematical modeling, we demonstrate that the anchoring strategy affects both component expression and diffusion, which in turn impact recruitment strength, kinetics, and spatial dynamics. Compared to the commonly used C-terminal iLID fusion, fusion proteins with large N-terminal anchors show stronger local recruitment, slower diffusion of recruited components, and efficient recruitment over wider gene expression ranges. We also define guidelines for component expression regimes for optimal recruitment for both cell-wide and subcellular recruitment strategies. Our findings highlight key sources of imprecision within light-inducible dimer systems and provide tools that allow greater control of subcellular protein localization across diverse cell biological applications.

**Significance:** Optogenetic light-inducible dimer systems, such as iLID, offer the ability to examine cellular signaling networks on second timescales and micrometer spatial scales. Confined light stimulation can recruit proteins to subcellular regions of the plasma membrane, and local signaling effects can be observed. Here, we report alternative iLID fusion proteins that display stronger and more spatially confined membrane recruitment. We also define optogenetic component expression regimes for optimal recruitment and show that slow-diffusing iLID proteins allow more robust recruitment in cell populations with heterogenous expression. These tools should improve the spatiotemporal control and reproducibility of optogenetic protein recruitment to the plasma membrane.

## Introduction

To execute fundamental processes including division, migration, and cell-to-cell communication, cells use signaling networks to organize protein activities within subcellular regions ^1–5^. Non-linear network topologies that include fast-acting feedback and crosstalk connections shape the outputs of these networks in space and time ^6,7^. However, identifying these motifs and understanding their roles have been pervasive challenges in cell biology. Recently, optogenetic tools have provided new opportunities to dissect complex signaling mechanisms by permitting specific activation of a single protein species at precise times and subcellular locations ^8–12^. In this way, signaling motifs can be directly probed to understand the roles of individual components and how they fit together to control complex cell behaviors.

The plasma membrane is a primary scaffold upon which signaling networks are organized ^13^. Indeed, concentrating upstream regulatory proteins, enzymes, and downstream effectors in distinct locations at the plasma membrane is a critical way to connect ligand sensing with localized signaling outputs. For instance, activated G-protein coupled receptors trigger asymmetric recruitment of lipid modifying enzymes and Rho GTPase regulators to polarize cells during migration ^14–16^. Receptor tyrosine kinases similarly recruit SH2 domain-containing proteins to distinct membrane locations to initiate diverse adhesive, metabolic, and proliferative signaling programs ^17^. Further downstream, actin regulating proteins are commonly recruited and concentrated at localized sites on the membrane to give cells shape, build specialized structures, or generate forces for movement ^18,19^. Engineering exogenous regulation of protein localization and activation at the plasma membrane represents an important step in dissecting, and ultimately controlling, diverse cell behaviors.

Optogenetic light-inducible dimer systems are especially valuable tools for rapid recruitment of specific proteins to the plasma membrane to study their effects with high spatial resolution. Several systems have been developed from naturally occurring photosensory protein families including phytochromes, cryptochromes, and light-oxygen-voltage (LOV) domain proteins ^9–12^. These domains vary in size, activation mechanism, and absorption wavelength to confer different properties to each system ^20^. Among the optogenetic systems, the improved Light-Induced Dimer (iLID) system has emerged as a leading tool because of its ease of use, fast kinetics, and large fold-change in component protein association ^12^. iLID is an engineered protein composed of a blue-light-sensing LOV2 domain and a flanking C-terminal helix (termed the Jα helix) fused to seven residues from the *E. coli* SsrA peptide that can bind with high affinity to the *E. coli* adaptor protein SspB ^17^. In iLID’s dark state conformation, tight “docking” of the Jα helix to the LOV core sterically prohibits SsrA-SspB interaction. However, upon irradiation with blue light, conformational changes in the LOV domain lead to helix undocking and heterodimerization between iLID and SspB. Dimerization occurs within seconds and persists until iLID reverts to its dark state. The dimer half-life (~30 seconds) is shorter than most other optogenetic tools, but it can still limit temporal experimental control. By anchoring iLID to the inner leaflet of the plasma membrane and fusing SspB to protein domains of interest, the system has been used to selectively activate Rho GTPases for migration, modulate Erk signaling during *Drosophila* development, and generate pulling forces for mitotic spindle positioning ^21–23^.

For maximum control of protein localization, an ideal light-inducible dimer system should display minimal interaction in the dark, a large fold-change in recruitment, and exact localization to the stimulated region for appropriate timescales. In practice, however, precision is limited by the physical properties of cells and biomolecules, including protein expression and diffusion. Due to non-negligible dark state binding and laws of mass action kinetics, the SspB component of the iLID system shows basal membrane localization that scales with protein levels ^24^. As a result, relative expression levels must be considered to strike a fine balance between basal and light-induced recruitment. Variability in expression on a cell-by-cell or construct-dependent basis exacerbates this issue, negatively impacting biological reproducibility and consistency across research groups ^25^. Additionally, lateral diffusion of the iLID component in the membrane limits spatial control for recruitment, especially for applications that require subcellular localization for extended periods of time ^26^. iLID is typically targeted to the membrane via fusion to the C-terminal CAAX motif of KRAS, which includes a post-translational prenylation site ^27^. This small lipid anchor could permit diffusion at a rate of ~0.5-1.7 μm^2^/sec, meaning that an activated iLID molecule may diffuse roughly 7 μm away from its site of activation before reverting to its dark state ^28,29^. Ultimately, these sources of noise and variability pose major limitations to optogenetic dimerization systems including iLID that have not been collectively addressed.

To alleviate these challenges, we generated iLID proteins with membrane anchors of different sizes and fusion configurations, and we systematically compared their subcellular recruitment dynamics using quantitative imaging. We showed that N-terminal anchoring configurations allow stronger recruitment compared to the existing C-terminal version, and that larger membrane anchors display slower diffusion and greatly improved spatial confinement of recruited proteins. We used mathematical modeling to define optogenetic component expression regimes to achieve optimal recruitment with different iLID fusions across diverse applications. Furthermore, we demonstrated that our slow-diffusing iLID proteins exhibit stronger local recruitment over wider component expression ranges.

## Results

### A diverse set of membrane-anchored iLID fusion proteins

We theorized that the C-terminal CAAX motif that is typically used to anchor iLID to the plasma membrane may influence subcellular recruitment dynamics (Figure 1A). First, the anchor’s proximity to the SsrA peptide could negatively affect iLID’s binding availability in the lit state. Second, the anchor’s small size and large diffusion coefficient contribute to the rapid spread of activated iLID molecules away from sites of stimulation, leading to poor spatial confinement of recruited proteins over time. To address these issues, we designed three alternative iLID fusion proteins that we hypothesized would show stronger recruitment and tighter localization (Figure 1B). To test the consequences of relocating the lipid anchor away from the SsrA peptide, we fused 11 residues from Lyn kinase, which contain post-translational myristoylation and palmitoylation sites, to the N-terminus of iLID (Lyn11-iLID) ^30^. Additionally, to compare protein confinement when iLID is anchored with slow-diffusing protein domains, we fused either the four-pass transmembrane protein Stargazin or the seven-pass transmembrane protein ADRB2 to the N-terminus of iLID (Stargazin-iLID and ADBR2-iLID, respectively) ^31,32^. Stargazin has been used previously to localize another optogenetic construct, LOVpep, to the membrane ^33^. We also fused mTurquoise2 directly upstream of iLID in all constructs to allow imaging of iLID expression and localization concurrent with activation using a 445 nm laser (Figure 1C). Each construct was stably expressed in HEK-293T cells, and imaging by confocal microscopy confirmed that all iLID constructs were sharply localized to the plasma membrane, although ADRB2-iLID fluorescence was noticeably dimmer than the other constructs (Figure 1D).

**Fig. 1.**
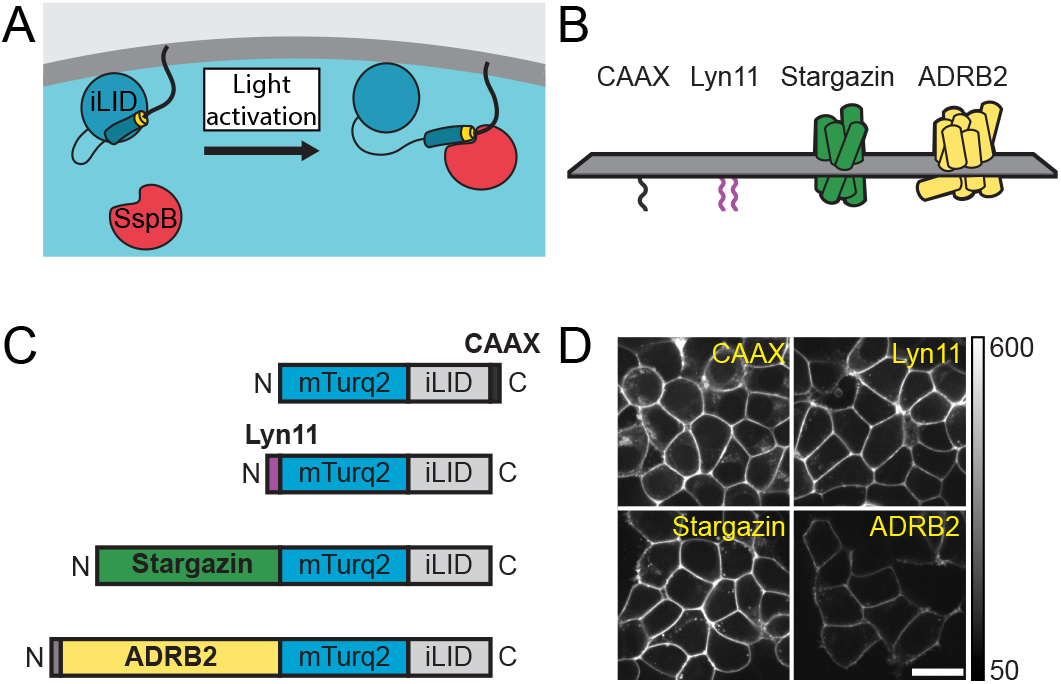
Overview of anchoring strategies for iLID fusion proteins. (A) Schematic of iLID-based membrane recruitment. The Jα helix is depicted as a blue-gray cylinder, the SsrA peptide is shown in yellow, and the Kras CAAX domain is depicted as a bold black line. (B) Illustration of anchors used to target iLID to the plasma membrane. The CAAX and Lyn11 sequences are membrane anchored via one and two lipid groups, respectively, while Stargazin and ADRB2 are four-pass and seven-pass transmembrane anchors, respectively. (C) Domain diagrams depicting the relative sizes and configurations of the iLID fusion proteins. (D) Comparison of localization of iLID constructs in HEK293T cells visualized by confocal microscopy. Imaging settings were kept consistent across constructs for easy comparison of relative brightness. Scale bar is 20 μm.

### N-terminal membrane anchors allow stronger recruitment

To assess recruitment magnitude and diffusive spread of the different iLID fusion proteins, we developed an automated point-recruitment assay using total internal reflection fluorescence (TIRF) microscopy. In TIRF microscopy, an induced evanescent field illuminates fluorophores exclusively within a ~100-200 nm thin region adjacent to the cell-glass interface. This imaging modality allowed us to monitor recruitment and diffusion of SspB molecules at the membrane while visualizing only a small fraction of background cytosolic protein (Supplemental Figure S1). We stably co-expressed each iLID construct with SspB fluorescently tagged with tandem dimer Tomato (tdTom-SspB) in HEK-293T cells. We monitored point-recruitment by performing TIRF imaging of tdTom-SspB every second for 25 seconds, with a single 10 ms pulse of 445 nm light applied within a ~3 μm radius spot at the cell centroid between frames 5 and 6 (Figure 2A). Importantly, we wrote a custom script to automate these experiments to reduce bias in cell selection, improve experimental reproducibility, and increase the number of biological replicates. To quantify recruitment magnitude, we calculated the fold change in SspB signal at the stimulus site by normalizing SspB values to the pre-stimulus level in each cell and calculating the mean time course over all cells. Intriguingly, Lyn11-iLID and Stargazin-iLID showed markedly higher recruitment compared to iLID-CAAX, while ADRB2-iLID showed slightly reduced recruitment (Figure 2B). Non-normalized SspB intensity before stimulation was similar for all conditions, though perhaps slightly elevated for Lyn11-iLID compared to other constructs. This suggested that differences in recruitment were primarily due to changes in binding interactions in iLID’s lit state (Figure 2C). It should be noted, however, that the non-normalized measurements depend on the degree of basal binding as well as total iLID and SspB protein levels in each cell. Therefore, we could not definitively assess possible differences in dark state binding between the iLID fusion proteins.

**Fig. 2.**
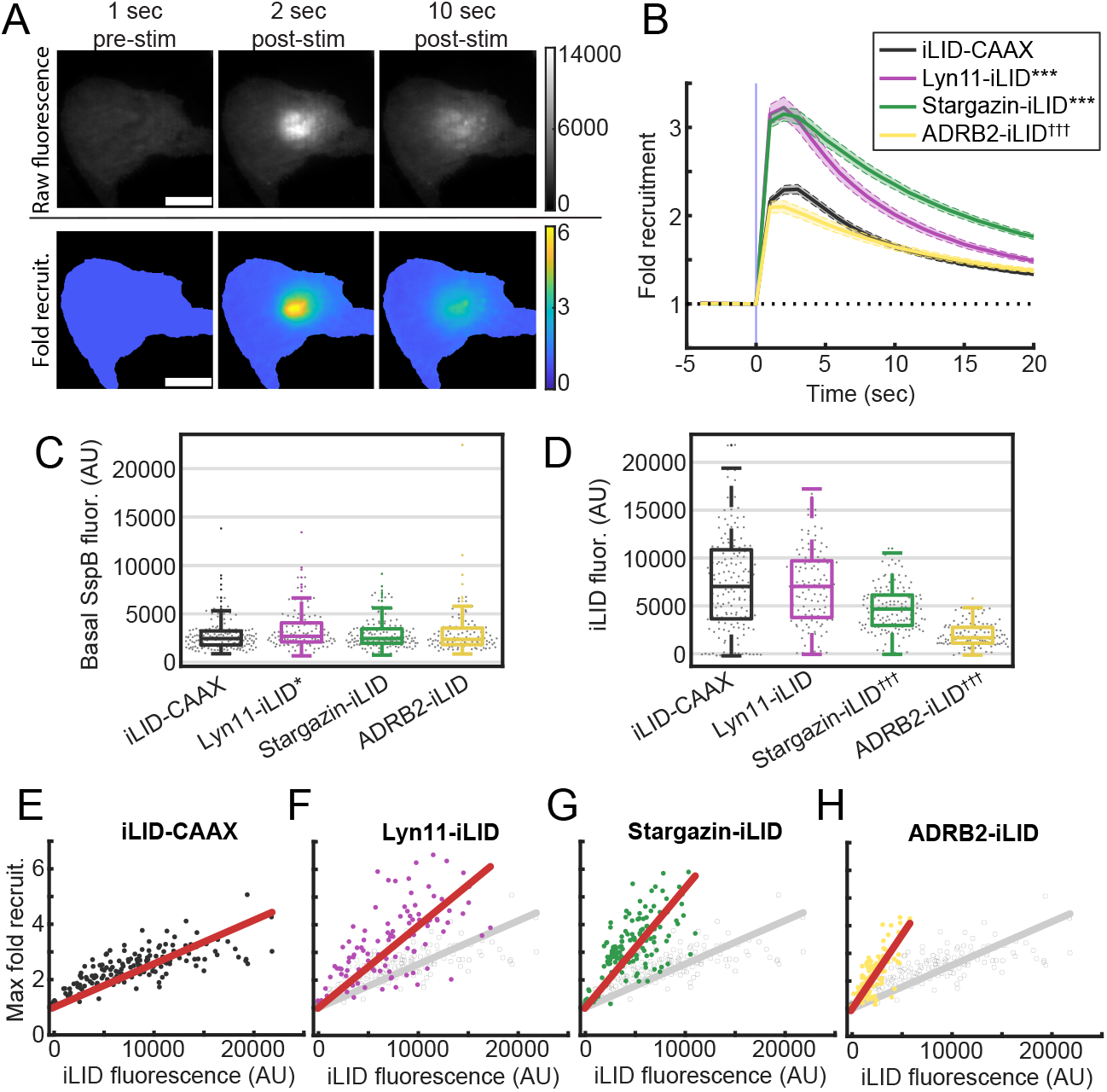
Analysis of recruitment magnitude via TIRF point-recruitment assay. (A) Example TIRF images of tdTom-SspB recruitment to the cell membrane before (left column) and after (middle and right columns) point-stimulation with a 10 ms pulse of 445 nm light. Top row shows raw fluorescence images and bottom row shows fold change in signal. Depicted cell expresses Lyn11-iLID. Scale bar is 15 μm. (B) Plots showing mean fold change in SspB signal at the stimulation site over time for each iLID construct, normalized to pre-stimulation levels. Error bars indicate ±S.E.M. (n=166 cells for iLID-CAAX, n=115 cells for Lyn11-iLID, n=144 cells for Stargazin-iLID, n=113 cells for ADRB2-iLID).***P<0.001 for maximum recruitment values above that of iLID-CAAX. ^†††^P<0.001 for maximum recruitment values below that of iLID-CAAX. P-values were calculated using the Mann-Whitney U test. (C-D) Box plots of average basal SspB fluorescence (C) or average iLID fluorescence (D) for each iLID construct. Central line indicates the median, bottom and top box edges indicate 25^th^ and 75^th^ percentiles, and whiskers indicate most extreme data points not considering outliers. Gray dots show distributions of single cell values (n=163 cells for iLID-CAAX, n=113 cells for Lyn11-iLID, n=143 cells for Stargazin-iLID, n=113 cells for ADRB2-iLID). *P<0.05 for values above that of iLID-CAAX. ^†††^P<0.001 for values below that of iLID-CAAX. P-values were calculated using the Mann-Whitney U test. (E-H). Scatterplots showing the correlation between iLID fluorescence and maximum fold recruitment in single cells. Red lines indicate the least-squares fit lines. For F-H, data from iLID-CAAX is shown in gray for easy comparison (n=163 cells for iLID-CAAX, n=113 cells for Lyn11-iLID, n=143 cells for Stargazin-iLID, n=113 cells for ADRB2-iLID).

Disparities in iLID expression levels between the constructs could account for the differences in SspB recruitment that we found. To investigate this possibility, we assessed relative iLID protein content at the membrane using fluorescence intensity from iLID TIRF images. We found that average fluorescence was highest for iLID-CAAX and Lyn11-iLID, ~33% lower for Stargazin-iLID, and ~75% lower for ADRB2-iLID (Figure 2D). Therefore, our alternative iLID fusion proteins show improved recruitment despite similar or lower expression levels. Notably, iLID-CAAX and Lyn11-iLID had roughly equivalent expression levels and differed only in the location of the lipid anchor, suggesting that the N-terminal location of the anchor is crucial for strong recruitment. In the case of ADRB2-iLID, drastically lower expression likely outweighs the potential improvements from N-terminal anchoring and results in a net reduction in SspB recruitment.

We observed considerable variability in iLID fluorescence at the single cell level, which we hypothesized could drive heterogeneity in SspB recruitment (Supplemental Figure S2A-S2D). Indeed, when we plotted the maximum fold change in SspB signal against iLID fluorescence in individual cells, we found a positive correlation for all constructs, supporting that higher iLID concentrations allow more SspB to be brought to the membrane upon stimulation (Figure 2E-H). Surprisingly, the slope of this correlation was larger for the N-terminally anchored iLID fusion proteins compared to iLID-CAAX. Therefore, at a given level of iLID expression, N-terminal anchors allow stronger SspB recruitment. Even more surprising, the slope of the correlation also increased corresponding to anchor size, suggesting that slow diffusion also contributes to higher local recruitment (Supplemental Figure S3). Taken together, our results demonstrate that localizing iLID to the membrane with a slow-diffusing N-terminal anchor enables stronger recruitment of SspB, even at lower iLID concentrations, and that this is achieved in large part through improved iLID-SspB interactions in the lit state.

### Mathematical models for guided optimization of iLID-based membrane recruitment

Since our point-recruitment assay results indicated that component expression has a major impact on recruitment efficiency, we developed differential equation models to define iLID and SspB expression regimes for optimal system performance. We modeled the iLID system as a reaction-diffusion system with four possible binding/activation states. Prior to stimulation, the system exists in a dynamic equilibrium between two states. In the first state, iLID molecules are inactive and unbound (free iLID_dark_) and SspB molecules are cytoplasmic (free SspB) (Figure 3A, top left). In the second state, SspB molecules are bound to inactive iLID at the membrane to form iLID_dark_-SspB dimers, which represents basal SspB recruitment (Figure 3A, bottom left). Upon light stimulation, iLID is rapidly converted to its active state, where it can then exchange between unbound (free iLID_lit_, free SspB) and bound (iLID_lit_-SspB) states (Figure 3A, top-right, bottomright). We assumed that activation and dark-state reversion of iLID occur independently of its binding state, with activation depending on the light intensity, light duration, and the iLID activation rate, and reversion depending on the intrinsic rate of iLID relaxation. Importantly, we were able to use previously reported measurements for iLID-SspB binding affinities in the lit (0.13 μM) and dark (4.7 μM) states and estimates of the iLID dark-state reversion rate (~0.02 s^−1^) to limit the number of free parameters in the model. Additionally, for modeling subcellular recruitment, we modeled the diffusion of all iLID-containing complexes with a single diffusion coefficient. We used the model to simulate the concentrations of its five molecular species (free iLID_dark_, free iLID_lit_, iLID_dark_-SspB, iLID_lit_-SspB, and free SspB) over the course of prototypical light-stimulation experiments.

**Fig. 3.**
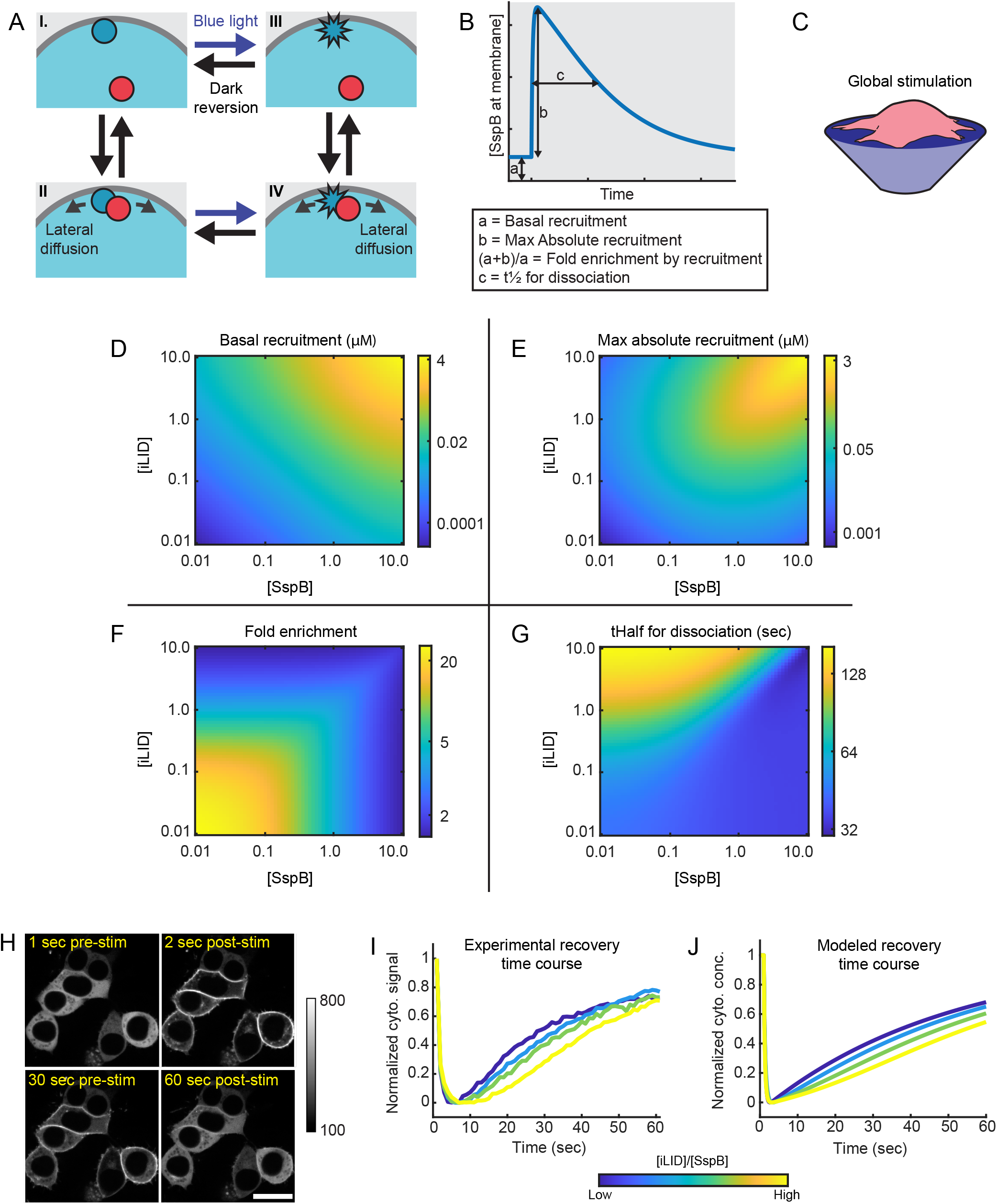
Overview of optimal expression regimes for global recruitment. (A) Schematic diagram illustrating the possible binding and activation states of the iLID system. Inactive iLID is represented as a blue circle, active iLID is represented as a blue 10-pointed polygon, and SspB is represented as a red circle. (B) Example plot of SspB concentration at the membrane over the course of a modeled light-stimulation experiment. Basal recruitment, max absolute recruitment, fold enrichment, and t_1/2_ for dissociation are shown based on predicted species concentrations. (C) Illustration of cell-wide iLID stimulation. (D-G) Heatmaps depicting extent of basal recruitment (D), max absolute recruitment (E), max fold enrichment by recruitment (F), and t_1/2_ for dissociation (G) across different iLID-to-SspB expression ratios generated using our ODE model. (H) Example confocal images showing distribution of SspB signal before (top-left panel) and after (top-right and bottom panels) global iLID stimulation using a 50 ms pulse of 445 nm light. Depicted cells express the Lyn11-iLID construct. Scale bar is 20 μm. (I) Plots of mean cytoplasmic SspB signal during confocal microscopy global stimulation experiments, with cells grouped based on estimated iLID-to-SspB ratio. The light pulse was applied between 1 and 2 seconds. Values are adjusted between 0 and 1 (From lowest to highest estimated iLID-to-SspB ratio: n=62 cells, n=54 cells, n=30 cells, n=55 cells). (J) For the experiment shown in H and I, predicted values were calculated from our ODE model using different iLID-to-SspB ratios and adjusted between 0 and 1. Total SspB concentration was 1.5 μM, and iLID concentrations were 50 nM (dark blue line), 1.5 μM (light blue line), 2 μM (green line), and 2.5 μM (yellow line).

### An optimal expression regime for cell-wide recruitment

Since iLID is often used to recruit proteins uniformly to the entire plasma membrane, we reasoned that global recruitment experiments provided both an ideal simplified system for validating our model, and a highly relevant system for optimizing component expression levels. Therefore, we first used an ordinary differential equation (ODE) version of our model lacking the diffusion terms to predict basal recruitment, maximum absolute recruitment, fold enrichment, and dissociation half-time upon cell-wide iLID activation (Figure 3B–3C). Note that, because TIRF measurements include signal from a small fraction of cytoplasmic protein (Supplemental Figure S1), empirical measurements of fold recruitment will differ from the predicted fold enrichment values, particularly for low iLID concentrations. We ran our model for a range of iLID-to-SspB ratios, titrating each component concentration between 10 nM and 10 μM to encompass the iLID-SspB affinity range for iLID dark and lit states.

As expected, both basal and max recruitment scaled with protein levels. At low iLID and SspB concentrations, our model showed minimal basal recruitment but weak absolute recruitment. In contrast, high iLID and SspB concentrations yielded high basal recruitment and strong absolute recruitment (Figure 3D-E). Because basal and absolute recruitment were correlated but with shifted dose-response curves, low component concentrations gave larger foldchanges in SspB recruitment (Figure 3F). Therefore, fold enrichment and absolute recruitment were largely inversely correlated. Notably, for a given level of basal recruitment, the maximum fold enrichment always occurred when iLID and SspB protein levels were equal (Figure 3D,F). Thus, for global recruitment strategies, equal expression of the two components is ideal, although the optimal levels will depend on a compromise between basal and stimulus-induced recruitment.

Interestingly, the model also made a less intuitive prediction: the time for iLID-SspB dissociation will be longer when iLID concentrations are high and in excess over SspB (Figure 3G). This effect is due to the increased likelihood of SspB re-binding to a neighboring molecule when more iLID is present at the membrane. We set out to test this prediction to validate our model, and to determine whether this effect occurs in the range of expression levels achieved in our cell lines. Using time lapse confocal microscopy, we measured the depletion and recovery of SspB signal in the cytoplasm before and after global iLID stimulation (Figure 3H). We observed characteristic rapid depletion then gradual recovery of cytoplasmic signal, though the magnitude and kinetics varied from cell to cell (Supplemental Figure S4). We organized the single cell data into four groups based on estimated iLID-to-SspB ratio and plotted curves for each group adjusted to a 0 to 1 scale to facilitate temporal comparisons. We used pre-stimulation SspB fluorescence as a measure of relative SspB concentration. Because iLID fluorescence in individual cells could not be measured directly in our assay, we used maximum percent signal loss to infer relative iLID concentrations, as iLID concentration and fold recruitment were strongly correlated in our previous experiments. In agreement with the model prediction, groups with higher estimated iLID-to-SspB ratios showed slower recovery of SspB signal and had recovery curves that qualitatively matched curves generated using our model (Figure 3I–3J).

Given the above findings, we suggest guidelines for component expression in optogenetic experiments using global recruitment. Low component concentrations (ideally around 60 nM) are preferable when fast kinetics are important and only low levels of SspB recruitment are necessary. For applications requiring higher levels of SspB recruitment, intermediate component concentrations (ideally in the 150 – 400 nM range) are optimal, as higher concentrations may suffer from high basal binding and poor enrichment. In both cases, the two component concentrations should be matched as closely as possible.

### An optimal expression regime for subcellular recruitment

The requirements for effective local recruitment can differ markedly from those for cell-wide recruitment. For instance, limitations on recruitment magnitude and kinetics may be altered since fewer component proteins participate, and the effect of diffusion becomes a major consideration. We therefore used a partial differential equation (PDE) model to assess optimal expression regimes for subcellular recruitment and investigate the effects of anchor diffusion (Figure 4A). As with our ODE model, we titrated iLID and SspB concentrations between 10 nM and 10 μM and calculated the expected fold enrichment by recruitment. We began by using an iLID diffusion coefficient of 1 μm^2^/sec as an estimate for the diffusion of iLID-CAAX, which falls within the range of measured diffusion rates for similar proteins ^28,29^. Distinct from our ODE model prediction, we found that optimal fold enrichment was achieved with iLID concentrations in excess of SspB by roughly 4-fold (Figure 4B). This arises because basal recruitment was similar to our ODE model predictions, while max absolute recruitment was high at lower SspB concentrations relative to iLID (Supplemental Figure S5A S5C). From these observations, we hypothesized that cells with high SspB concentrations would show lower fold recruitment on our point-recruitment assay. To investigate this, we overlaid single cell pre-stimulation SspB fluorescence values onto the correlation plots from Figure 2E–2H and looked for trends in recruitment according to basal SspB concentrations. It was important to compare only cells within conditions at similar iLID levels since pre-stimulation SspB fluorescence could also be affected by basal binding to the iLID components. Consistent with our expectations, we found that, at a given iLID concentration, cells with high SspB levels tended to have lower fold recruitment while cells with low SspB levels had higher fold recruitment (Supplemental Figure S6).

**Fig. 4.**
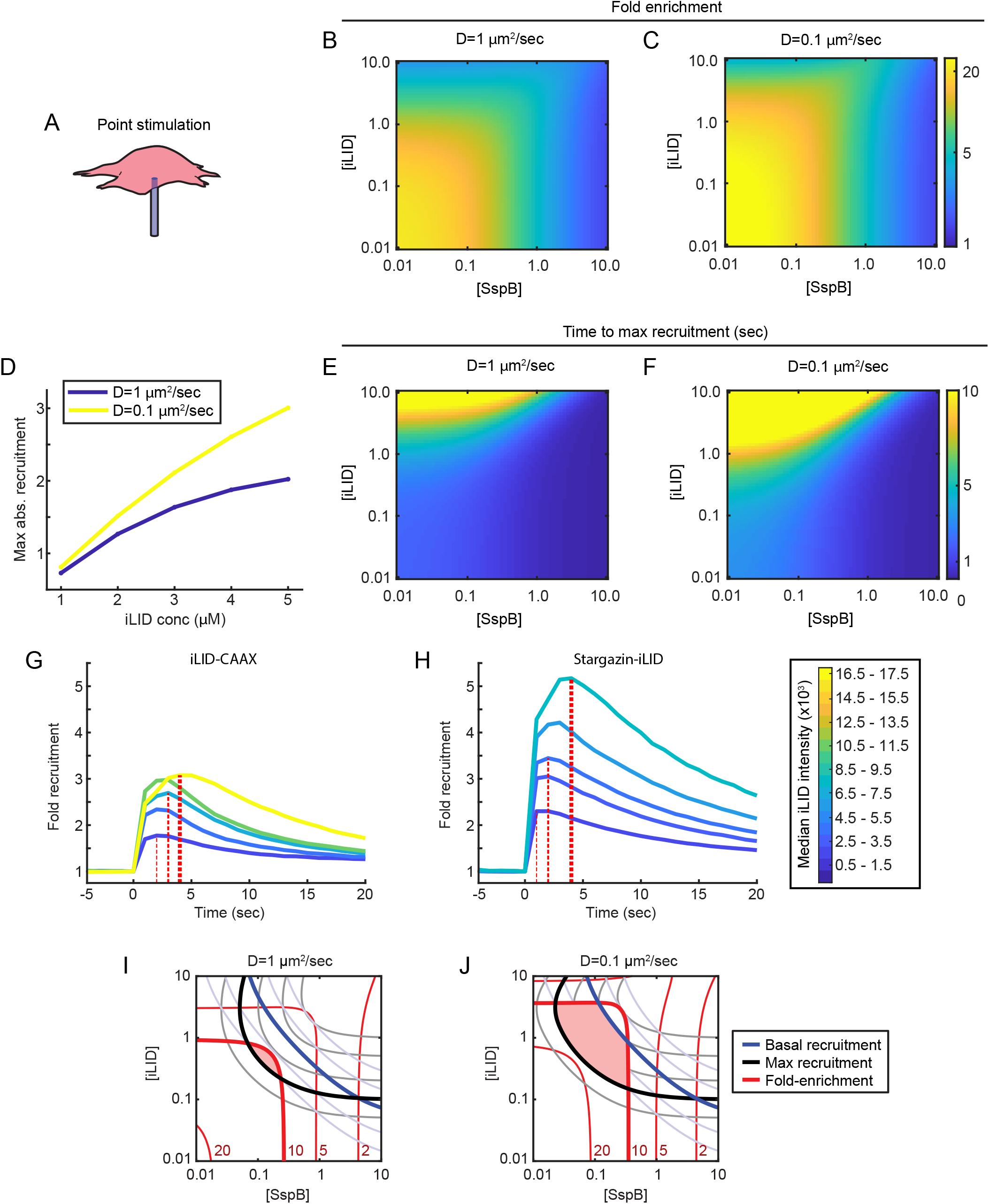
Overview of optimal expression regimes for subcellular recruitment. (A) Illustration of subcellular iLID stimulation. (B-C) Heatmaps depicting max fold enrichment by recruitment at the stimulation site across different iLID-to-SspB expression ratios generated using our PDE model. Values were generated using a diffusion coefficient of 1 μm^2^/sec (B) as an estimate for iLID-CAAX or 0.1 μm^2^/sec (C) as an estimate for Stargazin-iLID. (D) Plots of predicted maximum absolute recruitment at varying iLID concentrations using a diffusion coefficient of 1 μm^2^/sec (blue line) or 0.1 μm^2^/sec (yellow line). The total concentration of SspB was 1 μM. (E-F) Heatmaps depicting time to max recruitment across different iLID-to-SspB expression ratios generated using our PDE model. Values were generated using a diffusion coefficient of 1 μm^2^/sec (E) or 0.1 μm^2^/sec (F). (G-H) Plots showing fold recruitment at the stimulation site over time for cells grouped by relative iLID expression, calculated from cells expressing iLID-CAAX (G) or Stargazin-iLID (H). The line colors correspond to median iLID fluorescence intensities. T_max_ values for the first, third, and fifth groups are shown by thin, medium, and thick vertical lines, respectively (For iLID-CAAX, from lowest to highest iLID-to-SspB ratio: n=14 cells, n=13 cells, n=20 cells, n=25 cells, n=7 cells. For Stargazin-iLID, from lowest to highest iLID-to-SspB ratio: n=20 cells, n=14 cells, n=26 cells, n=12 cells, n=7 cells). (I-J) Overlaid contour plots of basal recruitment (blue lines), max recruitment (black lines), and fold enrichment by recruitment (red lines). Lines indicating 50 nM basal recruitment, 100 nM max recruitment, and 10-fold enrichment are boldened, and fold enrichment contour lines are labeled in red along the X-axis. The shaded area indicates component expression ranges predicted to give optimal recruitment. Plots were generated using our PDE model with a diffusion coefficient of 1 μm^2^/sec (I) or 0.1 μm^2^/sec (J).

Next, we asked whether the iLID diffusion coefficient affects the optimal component concentration regime. For comparison, we adjusted the iLID diffusion coefficient to 0.1 μm^2^/sec, an estimate for Stargazin-iLID based on diffusion rates of other transmembrane proteins ^34^. Intriguingly, we found that optimal recruitment was achieved with an even stronger asymmetry in concentrations, with up to almost 10-fold excess of iLID over SspB (Figure 4C). Again, this was due to high max absolute recruitment at even lower SspB concentrations (Supplemental Figure S5B and S5D). We hypothesized that, with slower diffusion, iLID molecules are better retained at the stimulation spot as maximum recruitment is being achieved, allowing higher absolute recruitment at low SspB levels. This would cause lower recruitment for fast-diffusing proteins compared to slow-diffusing proteins. To test this, we plotted the expected absolute recruitment for systems with diffusion coefficients of 1 or 0.1 μm^2^/sec. Across a range of iLID concentrations, modeled recruitment was higher with the slower diffusion coefficient (Figure 4D). Importantly, this finding provides an explanation for our previous observation that the slopes of the correlations between iLID fluorescence and recruitment increased with anchor size (Supplemental Figure S3).

Another prediction of our model is that the time to maximum recruitment (t_max_) depends on the membrane anchor diffusion and the relative component concentrations. We calculated t_max_ values at different iLID-to-SspB ratios with diffusion coefficients of 1 μm^2^/sec or 0.1 μm^2^/sec. Our model generally predicted higher t_max_ values with the slow-diffusing anchor (Figure 4E–4F). Interestingly, we also observed that t_max_ was markedly higher when iLID was in strong excess over SspB. This effect arises because iLID can deplete free SspB locally, and further recruitment depends on long-range SspB diffusion. To determine whether this behavior was evident in our point-recruitment experiments, we took cells with a narrow range of SspB fluorescence values, grouped them based on iLID fluorescence, and plotted average recruitment curves. For both iLID-CAAX and Stargazin-iLID cells, we observed the predicted shift in t_max_ values, and Stargazin-iLID showed shifted t_max_ values and higher recruitment magnitudes at lower iLID levels (Figure 4G–4H).

Ultimately, the iLID and SspB expression ranges for optimal subcellular recruitment depend on the nature of the iLID membrane anchor and the experimental goals. To explore this in more detail, we set out to characterize the conditions for optimal recruitment with both fast and slow diffusing anchors, given a set of desired system constraints. As a model case, we defined optimal recruitment as having no more than 50 nM basal recruitment, at least 100 nM max absolute recruitment, and at least a 10-fold increase in recruitment after stimulation. We then plotted contour lines for these three features and shaded the region in which all criteria were met. With fast diffusion, optimal recruitment was achieved within a relatively narrow range of expression levels (iLID concentrations between ~300 – 900 nM and SspB concentrations between ~75 – 250 nM) (Figure 4I). With slow diffusion, however, the range of iLID and SspB expression values for optimal recruitment was markedly wider, encompassing both higher and lower expression levels for each component (iLID concentrations between ~200 nM – 5 uM and SspB concentration between ~30 – 600 nM) (Figure 4J). This illustrates that experiments using slow-diffusing versions of iLID can better tolerate heterogeneity in expression levels, which is another key benefit to our alternative iLID fusions.

### Larger iLID anchors improve spatial confinement of recruitment

Our primary goal in building iLID membrane anchors with slower diffusion was to improve the spatial confinement of recruited proteins over time. To test whether this indeed occurred, we used the data from our point-recruitment assay to generate spatial SspB intensity gradients at each time point after stimulation. After normalizing SspB values to pre-stimulation levels, we measured the mean intensity as a function of radial distance from the stimulation site. Consistent with our expectations, we found that larger iLID anchor sizes resulted in improved confinement of SspB signal with sharper boundaries. As expected, the spatial extent of the initial SspB gradients right after stimulation was very similar for all iLID fusion proteins, despite differences in recruitment magnitude (Figure 5A–5D, dark green curves). However, at longer times after stimulation, SspB gradients spread and flattened to different extents that corresponded to the diffusion properties of the iLID membrane anchors. Signal spreading was most pronounced with iLID-CAAX, and only slightly improved with Lyn11-iLID, likely owing to the presence of an additional post-translational lipidation site (Figure 5A–5B, yellow curves, and Supplemental Movie S1 and S2). However, it was markedly improved for Stargazin-iLID and ADRB2-iLID (Figure 5C–5D, yellow curves, and Supplemental Movie S3 and S4). Indeed, when we used the distance for half-maximal gradient signal as a measure of spatial spread, we observed that between 1 and 10 seconds post-stimulation, the spread increased 65% for iLID-CAAX and 55% for Lyn11-iLID, but only 21% for Stargazin-iLID and 19% for ADRB2-iLID (Figure 5E).

**Fig. 5.**
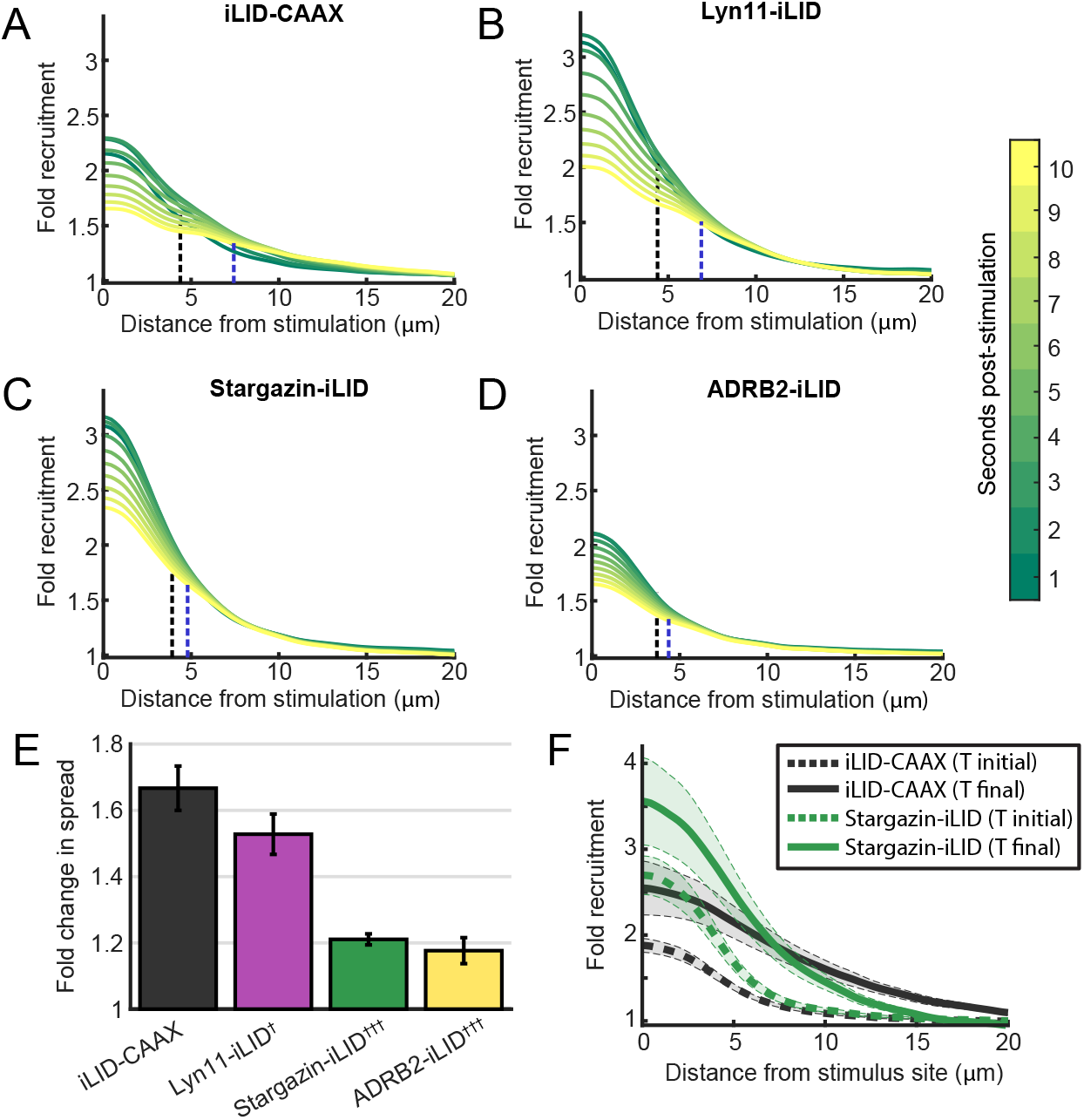
Spatial analysis of subcellular recruitment dynamics. (A-D) Plots depicting mean SspB intensity as a function of radial distance from the stimulation site. Values are normalized to pre-stimulation levels. Black and blue dotted lines indicate the distance of half-maximal gradient signal 1 second and 10 seconds after stimulation, respectively (n=156 cells for iLID-CAAX, n=110 cells for Lyn11-iLID, n=138 cells for Stargazin-iLID, n=107 cells for ADRB2-iLD). (E) Bar graphs showing fold change in distance of the half-maximal gradient signal from 1 to 10 seconds poststimulation, corresponding to the black and blue dotted lines in A-D. Error bars indicate ±S.E.M. ^†^P<0.05 and ^†††^P<0.001 for spread values below that of iLID-CAAX. P-values were calculated using the Mann-Whitney U test. (F) Plots of mean SspB intensity as a function of radial distance from the stimulation site after repeated point-stimulation for 1 minute. Dotted lines show intensity gradients 1 second after the initial stimulation and solid lines show intensity gradients 1 second after the final stimulation. iLID-CAAX and Stargazin-iLID gradients are overlaid for comparison of magnitude and spatial confinement. Error bars indicate ±S.E.M (n=24 cells for iLID-CAAX, n=22 cells for Stargazin-iLID).

Slow iLID diffusion is especially important to improve spatial confinement of recruitment over longer timescales. To more clearly demonstrate this, we performed a longer multi-stimulation recruitment experiment in which we repeatedly stimulated iLID every 10 seconds with a 10 ms light pulse over a 1-minute period. We then compared initial and final SspB gradients. Because Stargazin-iLID stood out as our best performing iLID fusion protein for subcellular recruitment, we limited our comparison to iLID-CAAX and Stargazin-iLID. With iLID-CAAX, the SspB gradient was initially localized, though relatively small in magnitude. Over time, the gradient spread and flattened drastically, showing elevated signal as far as 20 μm from the stimulation site after 1 minute (Figure 5F, black curves, and Supplemental Figure S7). On the other hand, with Stargazin-iLID, we observed stronger localized recruitment that remained more sharply localized over time, with higher signal at the stimulation site and markedly reduced signal at distal regions (Figure 5F, green curves, and Supplemental Figure S7). Ultimately, our results demonstrate that Stargazin-iLID is ideally suited for subcellular protein recruitment, as the construct expresses well and the large N-terminal transmembrane anchor allows stronger recruitment and increases confinement of signal through slower lateral diffusion.

## Discussion

Light-inducible dimers are highly flexible systems in a growing molecular toolkit for perturbing signaling networks in ever-more precise ways. With their acute spatial and temporal control, local signaling motifs such as feedback loops can be directly probed ^35^. Additionally, by adjusting light intensity, the appropriate amount of protein can be recruited to best suit the signaling system under investigation. However, these benefits are diminished by variable protein expression and diffusion, which limit the ability to reliably recruit the desired amount of protein to an exact subcellular location. In this study, we present alternative iLID fusion proteins that display stronger recruitment with higher spatial confinement across a wider range of component expression levels when compared to the previously published version. In doing so, we relied upon a combination of quantitative imaging and mathematical modeling. We found that experimental and conceptual approaches were both needed to dissect the combined influences of protein fusion configuration, diffusion, and component expression levels on optogenetic system performance. The use of automation in our live cell imaging approaches was critical in maximizing biological replicates and improving experimental reproducibility. This allowed us to assess each system quantitatively and draw direct comparisons to our mathematical model predictions.

Our alternative iLID fusion proteins offer a simple and adaptable way to improve local protein recruitment. Indeed, a key benefit to our approach is that different constructs can easily be interchanged depending on the specific requirements for a given experiment without introducing unintended biological effects. We recognize that our fusion proteins do not eliminate the issue of diffusion altogether, and other strategies have been put forward. For instance, efforts to engineer iLID and SspB variants with higher dissociation rate constants could improve spatial confinement, and one mutant protein with an altered dimer lifetime has been described ^24^. However, with fast dissociation, the total amount of recruited protein is diminished, and some downstream signaling processes that require longer activation times may not be triggered ^26,36^. Alternatively, Van Geel et al. restricted iLID diffusion directly using Cry2-based clustering of C2 domains or simultaneous anchoring to both membrane and microtubules ^37^. Their strategies greatly reduced iLID mobility through the membrane but come with their own important caveats. Heterogeneity in Cry2 expression and non-uniformity in microtubule positioning could diminish experimental reproducibility within their system, and intentional clustering of molecules and perturbations to microtubule dynamics could also alter biological outputs. Ultimately, the simplicity and overall effectiveness of our anchoring strategies make them an ideal option for subcellular optogenetic applications.

Aside from slowed diffusion, we observed a clear increase in fold recruitment when we switched from a C-terminal to N-terminal anchoring strategies. Based on our observations from TIRF microscopy, we hypothesize that the C-terminal CAAX anchor may lead to decreased flexibility or steric hindrance that limits interactions between SspB and the SsrA peptide within the Jα helix on iLID. If this is the case, it is possible that the CAAX anchor also plays a beneficial role in helping to prohibit dark-state iLID-SspB binding, which our N-terminal anchoring strategies may eliminate. While our TIRF data does not allow us to directly assess the extent of basal iLID-SspB binding, it does suggest that, on average, similar amounts of basal SspB are observed proximal to the cell membrane for all constructs. iLID’s LOV2 domain also contains a conserved N-terminal helix-turn-helix motif that engages in direct intramolecular interactions involved in photoswitching ^38^. Whether our N-terminal anchors affect these interactions is also unknown. In the future, determining the structural consequences of the various anchoring strategies may guide efforts to engineer iLID variants with even more precise photoswitching properties.

An unexpected benefit of our slow-diffusing iLID constructs is an increased expression range for which effective recruitment is still possible. A main difference between fast- and slow-diffusing proteins is a large increase in the range of iLID expression levels for which fold-change in recruitment remains high. Indeed, we frequently observed small fold changes in recruitment with the original iLID-CAAX system, which can likely be explained by the presence of only a small sub-population of cells with iLID expression levels that appropriately balance basal and light-activated recruitment. It is also worth noting that the comparisons we drew between fast- and slow-diffusing proteins with our model do not account for possible effects from using an N-terminal anchor. It is therefore possible that the true optimal expression ranges for Stargazin-iLID differ slightly from our model predictions. Finally, in all scenarios, we observed large changes in both basal and absolute recruitment over relatively small differences in component concentrations (Supplemental Figure S8). While our slow-diffusing constructs are more robust in cell populations with variable protein levels, care should be taken to minimize heterogenous expression by using stable rather than transient expression methods. Overall, our findings demonstrate the importance of considering the requirements for protein recruitment for different applications, and our model provides general guidelines for optimizing component expression patterns accordingly.

We expect that our alternative iLID fusion proteins will enable more precise subcellular recruitment and broaden the range of applications for which iLID can be used. Furthermore, the principles highlighted by our mathematical models will ensure greater reproducibility within and between research groups using iLID for light-based protein recruitment.

## Materials and Methods

### Cell culture

HEK-293T cells were obtained from the laboratory of Lifeng Xu and cultured in high glucose DMEM medium with sodium bicarbonate (Sigma-Aldrich catalog no D5671) and supplemented with 10% FBS, 1% penicillin and streptomycin, and 1% GlutaMax (Thermo Fisher catalog no 35050061) in a humidified incubator at 37° C and 5% atmospheric CO_2_. Cells were maintained at a density between 10% and 80% confluency on untreated 6-well tissue culture plates.

### Constructs

pLL7.0: Venus-iLID-CAAX (from KRas4B) and pLL7.0:tgRFPt-SSPB WT were gifts from Brian Kuhlman (Addgene plasmid #60411; http://n2t.net/addgene:60411; RRID:Addgene_60411 and Addgene plasmid #60415; http://n2t.net/addgene:60415; RRID:Addgene_60415) ^12^. Stargazin-GFP-LOVpep was a gift from Michael Glotzer (Addgene plasmid # 80406; http://n2t/net/addgene:80406; RRID:Addgene_80406) ^33^. Beta-2-adrenergic receptor-CFP was a gift from Catherine Berlot (Addgene plasmid # 55794; http://n2t.net/addgene:55794; RRID:Addgene_55794) ^39^. To generate the iLID plasmids, mTurqouise2, iLID-CAAX, iLID (no CAAX), the N-terminal myristoylated/palmitoylated domain of human Lyn tyrosine kinase (1-11), full-length Stargazin, and full-length ADRB2 were PCR amplified. Additionally, a modified version of the influenza hemagglutinin signal sequence (residues 2-16) was generated by oligonucleotide annealing for insertion upstream of the ADRB2 sequence to enhance membrane localization. Gene fragments were cloned via Gibson assembly into a lentiviral backbone containing an upstream ubiquitous chromatin opening element (UCOE), EF1-a promoter, and the blasticidin resistance gene downstream of an IRES sequence. For generating the tdTomato-SspB plasmid, td-Tomato, SspB, and iRFP-670 were PCR amplified. A tandem P2A/T2A sequence (tPT2A) reported by Liu et al. was synthesized using Twist Biosciences’ fragmentGene service ^40^. Gene fragments were cloned via Gibson assembly into a lentiviral backbone containing the EF1-a promoter and the puromycin resistance gene downstream of an IRES sequence. The downstream PT2A-iRFP-670 element allows for coexpression of iRFP-670 as a reference cytoplasmic marker if needed. Constructs were stably coexpressed in HEK-293T cells via subsequent rounds of lentivirus transduction and antibiotic selection.

### Automated TIRF Microscopy

Cells were plated on glass bottom 96-well plates (Cellvis catalog no P96-1.5H-N) treated with 200 ug/ml poly-D-lysine (Sigma-Aldrich catalog #P6407) and allowed to grow at 37° C and 5% atmospheric CO_2_ for ~16 hours. TIRF images were acquired using a Nikon Ti-E inverted fluorescence microscope with a 60x 1.49 NA Apo TIRF oil immersion objective and an Andor Zyla 4.2 sCMOS camera at 37° C. To minimize user bias and cell-to-cell variability, custom software was written for automated cell selection, cell stimulation and time lapse imaging using a MATLAB interface for Micro-manager. Briefly, each well was organized into an x-y matrix of locations that were imaged consecutively. At each location, a reference TIRF image was captured, and a correction was applied for non-uniform illumination of the TIRF field. From this, cell segmentation was performed using a minimum fluorescence threshold and minimum and maximum size thresholds. Cell masks were used to identify the cell with the highest area-to-perimeter ratio and move to its centroid coordinate. Area-to-perimeter ratio served as a suitable metric for selecting spread out cells with simple geometries. Finally, automated cell stimulation and time lapse imaging was carried out before moving to the next location. For single point-stimulation experiments, images were acquired every 1 second for 25 frames, and a single 10 ms pulse of 445 nm light was applied between frames 5 and 6. For multi-stimulation experiments, images were acquired every 2 seconds for 33 frames, and a 10 ms pulse of 445 nm light was applied after frames 3, 8, 13, 18, 23, 28, and 32. Subcellular iLID activation was achieved using a 445 nm laser connected to a Nikon FRAP illumination module with an ND 2.0 (1% transmission) neutral density filter in the light path. At the end of each experiment, a single TIRF image was acquired using a 445 nm laser to visualize iLID protein at the cell membrane.

### Determination of stimulus location for point-recruitment experiments

All imaging processing and analysis was performed using MATLAB. The stimulus location for point-recruitment experiments was estimated to be near the x-y center of the SspB fluorescence images, but the exact location could vary by a few pixels from day to day. Therefore, to determine the exact stimulus location for each day, the following empirical method was used. After image acquisition, cells with iLID fluorescence intensity below a minimum threshold, which was set manually based on background intensity, were excluded from analysis. Next, each movie was viewed in a randomized order and cells showing recruitment near the cell edge were eliminated. Remaining cells were used to generate a stimulus spot mask. For each cell, we performed a pixel-by-pixel subtraction of the raw 2 second post-stimulation SspB fluorescence image from the raw 1 second pre-stimulation SspB fluorescence image. The resulting images were composited, a rough stimulus spot mask was generated by masking for the brightest 20% of pixels, then a final stimulus spot mask was generated by finding the centroid coordinates of the rough mask and dilating the mask to a circle with a radius of 5 pixels (~1 μm).

### Background subtraction and cell segmentation of TIRF images

TIRF images were processed by performing local background subtraction and cell segmentation using previously described methods ^3^. Briefly, corrections were applied to raw fluorescence images to account for minor camera-specific artifacts and aberrations in the light-path. Background pixels were isolated by performing an initial conservative cell segmentation to remove high intensity pixels with fluorescence from cells or other objects. From the resulting background image, local background values were assigned by calculating the median intensity of pixels within defined 200×200 pixel blocks, then smoothing was applied to remove hard edges and generate a smooth background image. Next, cell masks were generated from the first SspB fluorescence image from each experiment. A single mask was suitable because no discernable changes to cell shape or location took place during individual experiments. To generate cell masks, smoothing was applied using a Gaussian filter of width 5 pixels (1.1 μm) to reduce pixel noise. Next, cell edges were enhanced using unsharp masking. Cell masks were generated using automated pixel thresholding followed by morphological closing (mask dilation followed by erosion), removal of small non-cell (<100 μm^2^) objects, and removal of masks of non-stimulated cells.

### Analysis of iLID and SspB intensity from TIRF images

Cells used to generate stimulus spot masks were included for analysis of iLID fluorescence intensity and SspB recruitment. To calculate iLID fluorescence intensities, background subtraction was performed as described above. iLID images were then computationally registered to SspB images using x and y translations. Images were smoothed with a Gaussian filter of width 2 pixels (0.32 μm) to reduce pixel noise and segmentation was performed by applying the previously generated cell masks. Finally, iLID fluorescence intensity was measured as the mean intensity of pixels within the stimulus spot mask. To calculate SspB florescence intensities, background subtraction and pixel smoothing were performed, and previously generated cell masks were applied. Basal SspB fluorescence intensity was measured as the mean intensity of pixels within the stimulus spot mask. For recruitment analysis, images were normalized to pre-stimulus levels by dividing each post-stimulation image by a reference 1 second pre-stimulation image on a pixel-by-pixel basis. SspB fluorescence intensity at the stimulation spot at each timepoint was calculated as the mean intensity of pixels within the stimulus spot mask. To calculate SspB signal for cell groups with different iLID levels (Figures 4G–4H), cells with basal SspB above 5,000 (AU) were eliminated to remove outliers. Cells were then binned into five groups based on iLID fluorescence intensity such that the magnitude of the recruitment curves were well-separated, and mean time courses were calculated for each group.

### Analysis of spatial SspB intensity gradients from TIRF images

The same images used for recruitment analysis were also used for spatial analysis, with the same image processing, smoothing, masking, and normalization applied. Using the stimulus spot masks and each cell mask, we first calculated the geodesic distance of each pixel in the cell mask from the site of stimulation, and grouped pixels based on distance values into bins of width 1.5 pixel (~0.32 μm). We then calculated the mean SspB intensity for each bin over time. The half-maximum of the gradient signal was calculated as the average of the maximum and minimum values of the gradient, and the distance for half-maximal signal was taken to be the distance of the gradient data point closest to the half-maximum value. Finally, spatial spread values were determined by calculating the fold change in half-maximum distance between 1 and 10 seconds post-stimulation (frames 6 and 16, respectively).

### Confocal Microscopy

Lyn11-iLID cells were used to study cytosolic SspB recovery via confocal microscopy. Cells were plated on glass bottom 96-well plates as described above. Confocal images were acquired using an Intelligent Imaging Innovations spinning disk confocal microscope using a 40x 1.3 NA Fluar oil immersion objective and Hamamatsu ORCA-Flash4.0 V2 sCMOS camera at 37° C. Time lapse imaging was conducted using 3i Slidebook6 software. For all experiments, images were acquired every 1 second. Visualization and photoactivation of iLID was performed by acquiring a single confocal image using a 445 nm laser.

### Analysis of cytoplasmic fluorescence intensity using confocal microscopy

Confocal time lapse image data was read into MATLAB using the Open Microscopy Environment’s Bio-Formats 6.1.0 toolbox. Time courses were analyzed in a randomized order to eliminate bias. For each time course experiment, we computed the fold change in SspB intensity from 1 second before to 3 seconds after stimulation on a pixel-by-pixel basis. Cytoplasmic areas of cells that responded to light stimulation were identified as large (>3.8 μm^2^) regions that showed at least 1% loss in pixel intensity over the chosen time frame. Next, a 25 pixel (~0.5 μm^2^) region of interest (ROI) was manually selected from each cytoplasmic area to ensure that cell edges and internal structures were not included in our analysis. Single cell intensity measurements were calculated as the mean SspB intensities of the ROIs at each time point. The non-normalized pre-stimulus SspB intensity was used to estimate total SspB abundance in each cell, and the percent loss in SspB intensity at 4 seconds post-stimulation (the time of peak recruitment for most cells) was used to estimate total iLID abundance. To assess recruitment and recovery kinetics at different component ratios, cells were binned into four groups of similarly sized ratio intervals, and average time courses were calculated.

### Mathematical modeling

All mathematical modeling was performed using code written in MATLAB. The objective of our differential equation model was to predict species concentrations over time at varying optogenetic component concentrations. Using the framework illustrated in Figure 3A, we identified five species that comprise the system: free iLID_dark_, free iLID_lit_, iLID_dark_-SspB, iLID_lit_-SspB, and free SspB. We began by defining parameters and rate constants for the activation and binding reactions based on published data. Reported equilibrium constants of 4.7 μM (Kd_dark_) and 130 nM (Kd_lit_) for iLID_dark_ and iLID_lit_, respectively, were used ^12^. The iLID-SspB association rate (R_assoc._) was assumed to be the same for both dark and lit states of iLID, the lit state dissociation rate was estimated to be of 0.5 sec^−1^, and an iLID dark-state reversion rate (R_rev._) of 0.02 sec^−1^ was chosen based on published measurements ^26^. For the PDE model, we also included a diffusion coefficient parameter that could be set according to the iLID construct being modeled and a cell radius parameter that was set to 15 μm. We used a diffusion coefficient of 25 μm^2^/sec for free SspB based on published estimates for cytosolic proteins ^41^. Other kinetic parameters could be defined from the above:

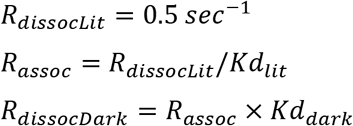

We next determined initial conditions, assuming a steady-state in which no iLID molecules were active before stimulation. Therefore, [*initial free iLID_lit_*] = 0 and [*initial iLIDiit – SspB*] = 0. We then used conservation of mass and equilibrium binding to compute the initial free iLID_dark_, initial free SspB, and initial iLID_dark_-SspB complex as follows:

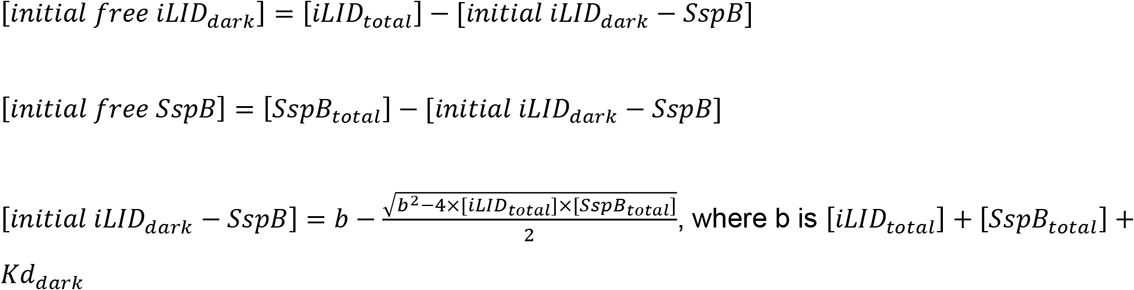

For the PDE model, we also defined symmetry and boundary conditions to account for spatial considerations. We modeled the cell as a radially symmetric disc with membrane and cytoplasmic compartments modeled at every radial distance from a center stimulation point to the outer cell edge, and we used a reflective boundary condition.

For the ODE model, we included a stepwise function as an input to our model to mimic global iLID stimulation, with a single stimulation pulse at the start of the model time course equivalent to 100 ms in length. We assumed that all iLID molecules were converted to their active form upon stimulation. For the PDE model, we mimicked subcellular light stimulation by generating an input function that was fit to match the spatial SspB intensity gradient 1 second after stimulation from Stargazin-iLID data, scaled to have values between 0 and 1. We refer to the solution of these input functions as Stim_fun_ in the equations below, and note that Stim_fun_ was defined to be uniformly zero at all times other than during simulated light stimulation.

Finally, we generated the following differential equations to define the concentration of each species:

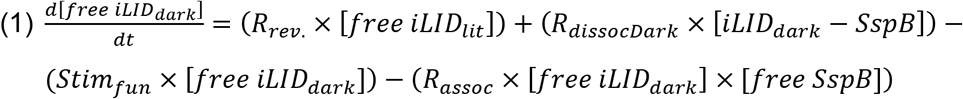

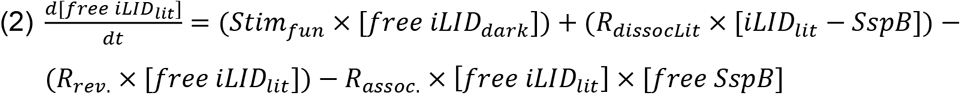

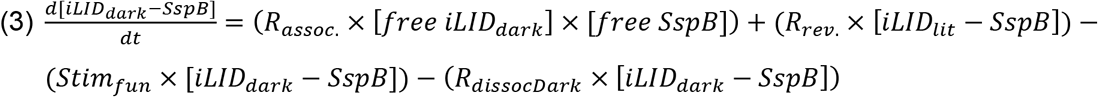

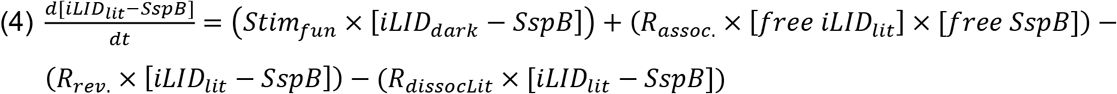

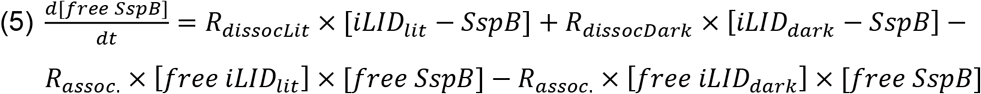

To run our model calculations, we used a first-order ODE solver (the ode45 function in MATLAB) and a 1-D PDE solver (the pdepe function in MATLAB) for the ODE and PDE versions of the model, respectively. A 1-D solver was appropriate for our PDE model given the radial symmetry of the system.

### Statistics

Statistical analyses were carried out in MATLAB. For Figure 2B–2D and Figure 5E, significance was calculated using the Mann-Whitney U test (ranksum function in MATLAB) to compare values from each construct to iLID-CAAX. All results in this study were independently replicated at least three times.

### Data availability

Raw datasets and analysis code are available upon request.

## Acknowledgements

We thank Dr. Michael Paddy and the UC Davis MCB Light Microscopy Imaging Facility for providing equipment and guidance for confocal microscopy experiments, and Annalise Gushue for assistance in cloning optogenetic constructs. We also thank George Bell, Briana Rocha-Gregg, Stefan Lundgren, Emel Akdogan, Diana Sernas, Samuel Hayes, Arthur Mercer, and Esther Rincón Gila for critical discussion and assessment of the manuscript. This work was funded by an NIH Director’s New Innovator Award (DP2 HD094656) to SRC. DEN was supported by an NIH T32 Fellowship (T32 GM007377) and an NIH F31 Fellowship (F31 HL152621-01).

## Author contributions

D.E.N. and S.R.C. conceived and designed the experiments. D.E.N. generated all constructs and cell lines, and performed all experiments and data analyses. S.R.C. wrote the mathematical models and oversaw their application by D.E.N. D.E.N. wrote the manuscript with input from S.R.C. and acknowledged colleagues.

## References

1. Yüce, Ö., Piekny, A. & Glotzer, M. An ECT2-centralspindlin complex regulates the localization and function of RhoA. J. Cell Biol. 170, 571–582 (2005).

2. Janetopoulos, C., Ma, L., Devreotes, P. N. & Iglesias, P. A. Chemoattractant-induced phosphatidylinositol 3,4,5-trisphosphate accumulation is spatially amplified and adapts, independent of the actin cytoskeleton. Proc. Natl. Acad. Sci. U. S. A. 101, 8951–8956 (2004).

3. Yang, H. W., Collins, S. R. & Meyer, T. Locally excitable Cdc42 signals steer cells during chemotaxis. Nat. Cell Biol. 18, 191–201 (2016).

4. Reinhard, N. R. et al. The balance between Gαi-Cdc42/Rac and Gα12/13 -RhoA pathways determines endothelial barrier regulation by sphingosine-1-phosphate. Mol. Biol. Cell 28, 3371–3382 (2017).

5. Singleton, K. L. et al. Spatiotemporal patterning during T cell activation is highly diverse. Sci. Signal. 2, 1–13 (2009).

6. Brandman, O. & Tobias. Feedback Loops Shape Cellular Signals in Space and Time. Science (80-.). 322, 390–395 (2008).

7. Guilluy, C., Garcia-Mata, R. & Burridge, K. Rho protein crosstalk: Another social network? Trends Cell Biol. 21, 718–726 (2011).

8. Yazawa, M., Sadaghiani, A. M., Hsueh, B. & Dolmetsch, R. E. Induction of protein-protein interactions in live cells using light. Nat. Biotechnol. 27, 941–945 (2009).

9. Levskaya, A., Weiner, O. D., Lim, W. A. & Voigt, C. A. Spatiotemporal control of cell signalling using a light-switchable protein interaction. Nature 461, 997–1001 (2009).

10. Kennedy, M. J. et al. Rapid blue-light-mediated induction of protein interactions in living cells. Nat. Methods 7, 973–977 (2010).

11. Strickland, D. et al. TULIPs: tunable, light-controlled interacting protein tags for cell biology. Nat. Methods 9, 379–84 (2012).

12. Guntas, G. et al. Engineering an improved light-induced dimer (iLID) for controlling the localization and activity of signaling proteins. Proc. Natl. Acad. Sci. 112, 112–117 (2015).

13. Grecco, H. E., Schmick, M. & Bastiaens, P. I. H. Signaling from the living plasma membrane. Cell 144, 897–909 (2011).

14. Parent, C. A., Blacklock, B. J., Froehlich, W. M., Murphy, D. B. & Devreotes, P. N. G protein signaling events are activated at the leading edge of chemotactic cells. Cell 95, 81–91 (1998).

15. Iijima, M. & Devreotes, P. Tumor Suppressor PTEN Mediates Sensing of Chemoattractant Gradients. Cell 109, 599–610 (2002).

16. Xu, J. et al. Divergent signals and cytoskeletal assemblies regulate self-organizing polarity in neutrophils. Cell 114, 201–214 (2003).

17. Dougan, D. A., Weber-ban, E., Bukau, B. & Heidelberg, B. Targeted Delivery of an ssrA-Tagged Substrate by the Adaptor Protein SspB to Its Cognate AAA+ Protein ClpX. Mol. Cell 12, 373–380 (2003).

18. Rafelski, S. M. & Theriot, J. A. Mechanism of polarization of Listeria monocytogenes surface protein ActA. Mol. Microbiol. 59, 1262–1279 (2006).

19. Weiner, O. D., Marganski, W. A., Wu, L. F., Altschuler, S. J. & Kirschner, M. W. An actin-based wave generator organizes cell motility. PLoS Biol. 5, 2053–2063 (2007).

20. Tischer, D. & Weiner, O. D. Illuminating cell signalling with optogenetic tools. Nat. Rev. Mol. Cell Biol. 15, 551–558 (2014).

21. O’Neill, P. R., Kalyanaraman, V. & Gautam, N. Subcellular optogenetic activation of Cdc42 controls local and distal signaling to drive immune cell migration. Mol. Biol. Cell 27, 1442–1450 (2016).

22. Johnson, H. E. et al. The Spatiotemporal Limits of Developmental Erk Signaling. Dev. Cell 40, 185–192 (2017).

23. Okumura, M., Natsume, T., Kanemaki, M. T. & Kiyomitsu, T. Dynein–dynactin–NuMA clusters generate cortical spindle-pulling forces as a multiarm ensemble. Elife 7, 1–24 (2018).

24. Zimmerman, S. P. et al. Tuning the Binding Affinities and Reversion Kinetics of a Light Inducible Dimer Allows Control of Transmembrane Protein Localization. Biochemistry 55, 5264–5271 (2016).

25. Toettcher, J. E., Gong, D., Lim, W. A. & Weiner, O. D. Light-based feedback for controlling intracellular signaling dynamics. Nat. Methods 8, 837–839 (2011).

26. Benedetti, L. et al. Light-activated protein interaction with high spatial subcellular confinement. Proc. Natl. Acad. Sci. U. S. A. 115, E2238–E2245 (2018).

27. Apolloni, A., Prior, I. A., Lindsay, M., Parton, R. G. & Hancock, J. F. H-ras but Not K-ras Traffics to the Plasma Membrane through the Exocytic Pathway. Mol. Cell. Biol. 20, 2475–2487 (2000).

28. Das, S. et al. Single-molecule tracking of small gtpase RAC1 uncovers spatial regulation of membrane translocation and mechanism for polarized signaling. Proc. Natl. Acad. Sci. U. S. A. 112, E267–E276 (2015).

29. Meder, D., Moreno, M. J., Verkade, P., Vaz, W. L. C. & Simons, K. Phase coexistence and connectivity in the apical membrane of polarized epithelial cells. Proc. Natl. Acad. Sci. U. S. A. 103, 329–334 (2006).

30. Kovářová, M. et al. Structure-Function Analysis of Lyn Kinase Association with Lipid Rafts and Initiation of Early Signaling Events after Fcε Receptor I Aggregation. Mol. Cell. Biol. 21, 8318–8328 (2001).

31. Letts, V. A. et al. The mouse stargazer gene encodes a neuronal Ca2+-channel γ subunit. Nat. Genet. 19, 340–347 (1998).

32. Bang, I. & Choi, H. J. Structural features of β2 adrenergic receptor: Crystal structures and beyond. Mol. Cells 38, 105–111 (2015).

33. Wagner, E. & Glotzer, M. Local RhoA activation induces cytokinetic furrows independent of spindle position and cell cycle stage. J. Cell Biol. 213, 641–649 (2016).

34. Yanagawa, M. et al. Single-molecule diffusion-based estimation of ligand effects on G protein-coupled receptors. Sci. Signal. 11, 1–16 (2018).

35. Lamas, I., Merlini, L., Vještica, A., Vincenzetti, V. & Martin, S. G. Optogenetics reveals Cdc42 local activation by scaffold-mediated positive feedback and Ras GTPase. PLoS Biology vol. 18 (2020).

36. Tummino, P. J. & Copeland, R. A. Residence time of receptor - Ligand complexes and its effect on biological function. Biochemistry 47, 5481–5492 (2008).

37. Van Geel, O., Hartsuiker, R. & Gadella, T. W. J. Increasing spatial resolution of photoregulated GTPases through immobilized peripheral membrane proteins. Small GTPases 00, 1–10 (2018).

38. Halavaty, A. S. & Moffat, K. N- and C-terminal flanking regions modulate light-induced signal transduction in the LOV2 domain of the blue light sensor phototropin 1 from Avena sativa. Biochemistry 46, 14001–14009 (2007).

39. Hynes, T. R., Mervine, S. M., Yost, E. A., Sabo, J. L. & Berlot, C. H. Live cell imaging of Gs and the β2-adrenergic receptor demonstrates that both αs and β 1γ7 internalize upon stimulation and exhibit similar trafficking patterns that differ from that of the β2-adrenergic receptor. J. Biol. Chem. 279, 44101–44112 (2004).

40. Liu, Z., Chen, O., Wall, J. B. J., Zheng, M. & Zhou, Y. Systematic comparison of 2A peptides for cloning multi-genes in a polycistronic vector. Sci. Rep. 7, 1–9 (2017).

41. Swaminathan, R., Hoang, C. P. & Verkman, A. S. Photobleaching recovery and anisotropy decay of green fluorescent protein GFP-S65T in solution and cells: Cytoplasmic viscosity probed by green fluorescent protein translational and rotational diffusion. Biophys. J. 72, 1900–1907 (1997).

