## Supplemental Figures and Legends for "Optimized iLID membrane anchors for local optogenetic protein recruitment"

Figure S1. Molecule-scale illustration showing how SspB recruitment is visualizated via TIRF microscopy. SspB molecules are represented as red circles and iLID molecules are represented as blue circles. When incident light hits the sample at a critical angle, total internal reflection creates an evanescent field that illuminates fluorescent proteins within 100-200 nm of the basal plasma membrane. Prior to stimulation, most SspB molecules are unbound from iLID and not visible by TIRF. However, a small fraction of cytoplasmic SspB is proximal to the membrane and visible. After stimulation, SspB molecules that bind to activated iLID are recruited into the field of view at high concentrations.

Figure S2. Histograms of maximum fold change in recruitment in individual cells from point-recruitment experiments. Histrograms show single cell data for (A) iLID-CAAX, (B) Lyn11-iLID, (C) Stargazin-iLID, and (D) ADRB2-iLID. Bin edges and widths were constant for all constructs. In B-D, iLID-CAAX data is overlaid as red stairs for easy comparison.

Figure S3. Fit lines from Figures 2E-2H are overlaid. The slope of the correlation is substantially steeper for N-terminal versus C-terminal anchors, and also increases according to anchor size.

Figure S4. Plots of cytoplasmic SspB signal in Lyn11-iLID expressing cells during confocal microscopy global stimulation experiments. Single cell traces are shown in gray, and mean trace of all cells is shown in red. The light pulse was applied between 1 and 2 seconds. Values are normalized to pre-stimulus levels such that recruitment and recovery are depicted as fractions of total SspB signal.

Figure S5. Recruitment heatmaps from PDE model predictions. (A-B) Heatmaps of basal recruitment at iLID and SspB concentrations ranging from 10 nM to 10 $\mu$M, using a diffusion coefficient of 1 $\mu m^2$/sec (A) or 0.1 $\mu m^2$/sec (B). Heatmaps are identical with both diffusion rates, since basal recruitment occurs uniformly throughout the membrane. (C-D) Heatmaps of max absolute recruitment at iLID and SspB concentrations ranging from 10 nM to 10 $\mu$M, using a diffusion coefficient of 1 $\mu m^2$/sec (C) or 0.1 $\mu m^2$/sec (D). Note the asymmetry in absolute recruitment with increasing component concentrations, which becomes more exaggerated with a smaller diffusion coefficient.

Figure S6. Correlation plots of iLID fluorescence vs max fold recruitment after point-stimulation for iLID-CAAX (A), Lyn11-iLID (B), Stargazin-iLID (C), and ADRB2-iLID (D), as shown in Figures

2E-2H. Single cell data points are pseudocolored according to basal SspB intensity, with low SspB intensity in pink and high SspB intensity in yellow. Pseudocolor ranges are set according to the minimum and maximum basal SspB intensity for each condition.

Figure S7. Spatial SspB intensity gradients for iLID-CAAX (black) and Stargazin-iLID (green) from multi-stimulation experiments. Dotted lines show intensity gradients 1 second after the initial stimulation and solid lines show intensity gradients 1 second after the final stimulation. Values are adjusted to a scale from 0 to 1, with 0 corresponding to the minimum value of initial gradients and 1 corresponding to the maximum value of final gradients. This representation corrects for differences in recruitment magnitude and allows easy comparison of the extent of signal spreading for each construct.

Figure S8. Visualization of ODE recruitment heatmaps as line plots. Plots of basal recruitment (A) and max absolute recruitment (B) as a function of iLID concentration, and generated across a range of SspB concentrations (depicted as different line colors). Component concentrations range from 10 nM to 10 μM. Both features show steep increases, especially at mid-range to high component concentrations.

Movie S1. Time lapse TIRF imaging of tdTom-SspB recruitment to the plasma membrane with the iLID-CAAX construct. A HEK-293T cell co-expressing iLID-CAAX and tdTom-SspB was locally stimulated with a 10ms pulse of 445 nm light at its center. After stimulation, tdTom-SspB accumulates at the stimulation site, then rapidly diffuses radially toward the cell periphery. Pixel intensity bounds are 0 – 19,000 (AU). Scale bar is 15 μm.

Movie S2. Time lapse TIRF imaging of tdTom-SspB recruitment to the plasma membrane with the Lyn11-iLID construct. A HEK-293T cell co-expressing Lyn11-iLID and tdTom-SspB was locally stimulated with a 10ms pulse of 445 nm light at its center. Diffusion of tdTom-SspB away from the stimulation site is still easily observable, despite the additional lipidation site on the Lyn11 fragment. Pixel intensity bounds are 0 – 12,000 (AU). Scale bar is 15 μm.

Movie S3. Time lapse TIRF imaging of tdTom-SspB recruitment to the plasma membrane with the Stargazin-iLID construct. A HEK-293T cell co-expressing Stargazin-iLID and tdTom-SspB was locally stimulated with a 10ms pulse of 445 nm light at its center. Compared to cells expressing iLID-CAAX and Lyn11-iLID, diffusion of tdTom-SspB is markedly reduced. Pixel intensity bounds are 0 – 17,500 (AU) Scale bar is 15 μm.

Movie S4. Time lapse TIRF imaging of tdTom-SspB recruitment to the plasma membrane with the ADRB2-iLID construct. A HEK-293T cell co-expressing ADRB2-iLID and tdTom-SspB was locally stimulated with a 10ms pulse of 445 nm light at its center. tdTom-SspB diffusion is the mostly highly spatially restricted in cells expressing ADRB2-iLID. Pixel intensity bounds are 0 – 14,000 (AU) Scale bar is 15 μm.

### Figure S1

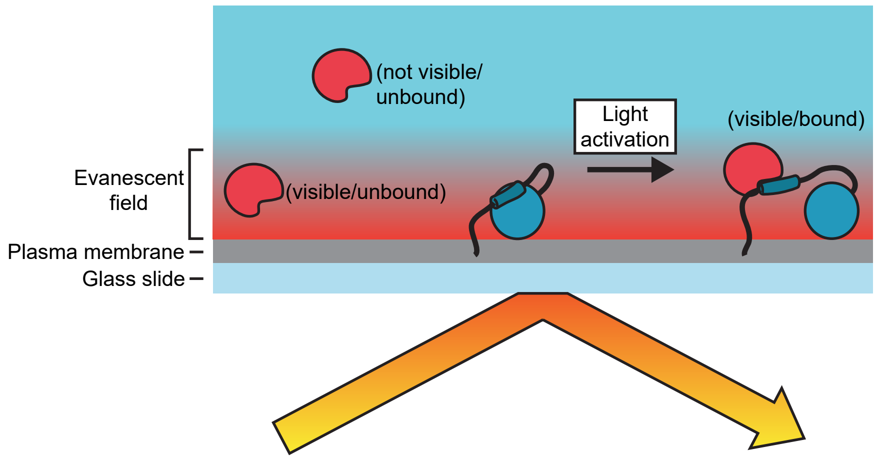

(not visible/
unbound)
Light
activation
(visible/bound)
Evanescent
field
(visible/unbound)
Plasma membrane
Glass slide

Figure S2

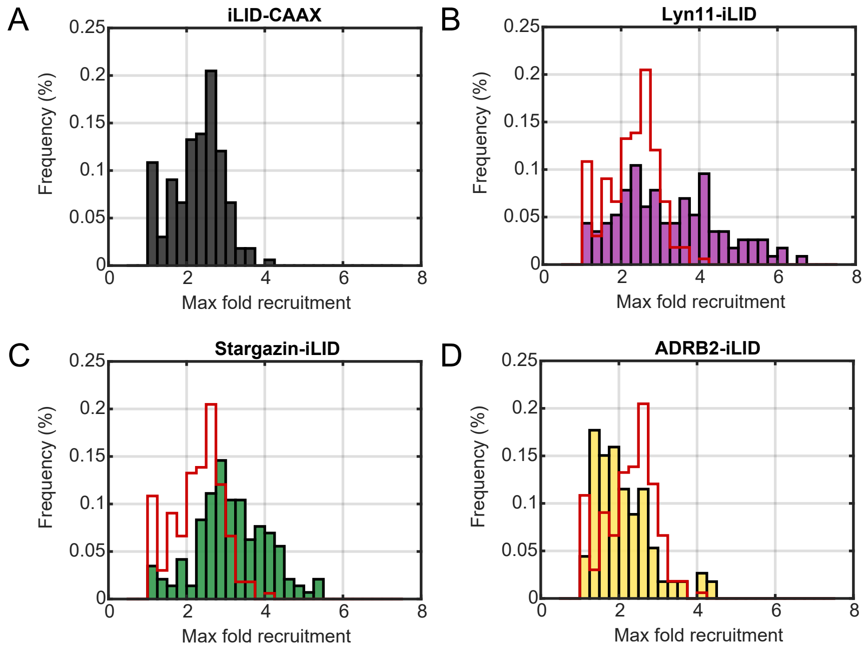

A
iLID-CAAX
B
Lyn11-iLID
C
Stargazin-iLID
D
ADRB2-iLID
Frequency (%)
Max fold recruitment
0.25
0.2
0.15
0.1
0.05
0
0
2
4
6
8

### Figure S3

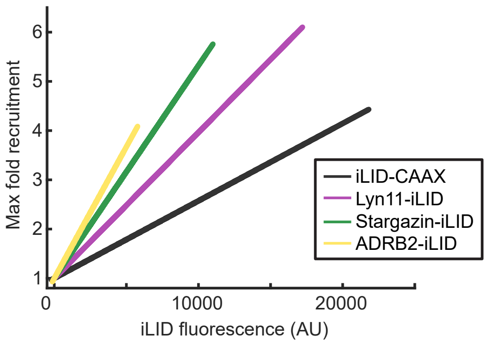

Max fold recruitment
1
2
3
4
5
6
0
10000
20000
iLID fluorescence (AU)
iLID-CAAX
Lyn11-iLID
Stargazin-iLID
ADRB2-iLID

### Figure S4

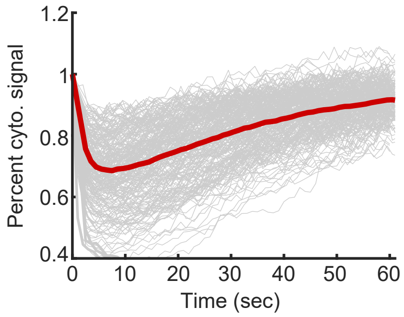

### Figure S5

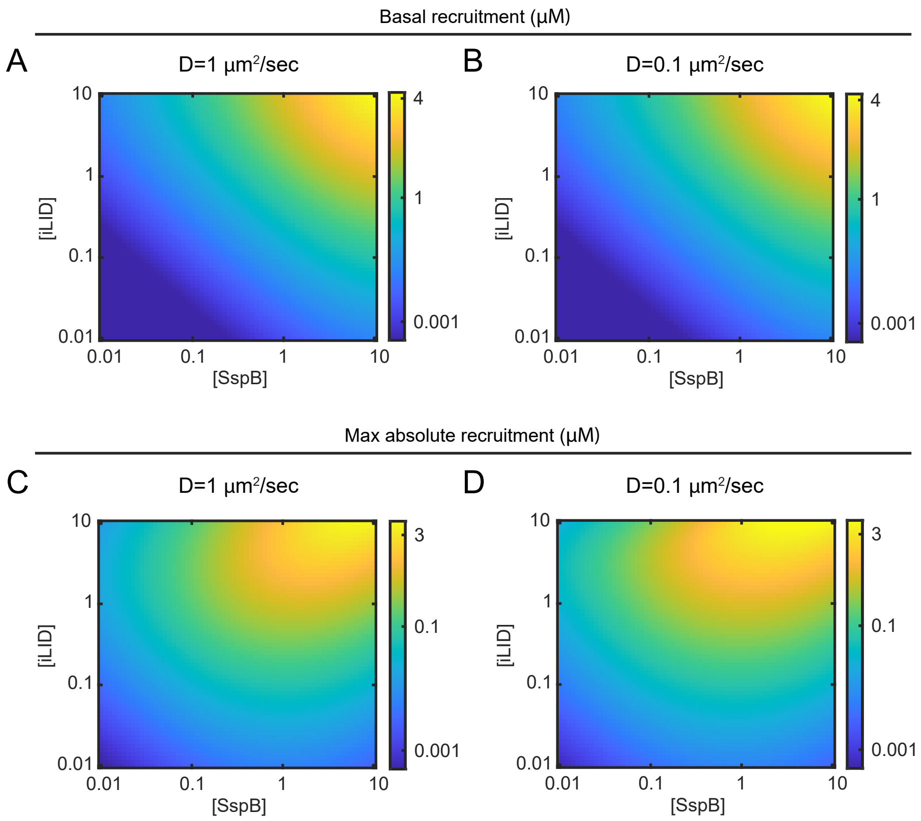
Basal recruitment (µM)
A
D=1 µm²/sec
B
D=0.1 µm²/sec
[iLID]
[SspB]
10
1
0.1
0.01
0.01
0.1
1
10
4
1
0.001
Max absolute recruitment (µM)
C
D=1 µm²/sec
D
D=0.1 µm²/sec
3
0.1
0.001

### Figure S6

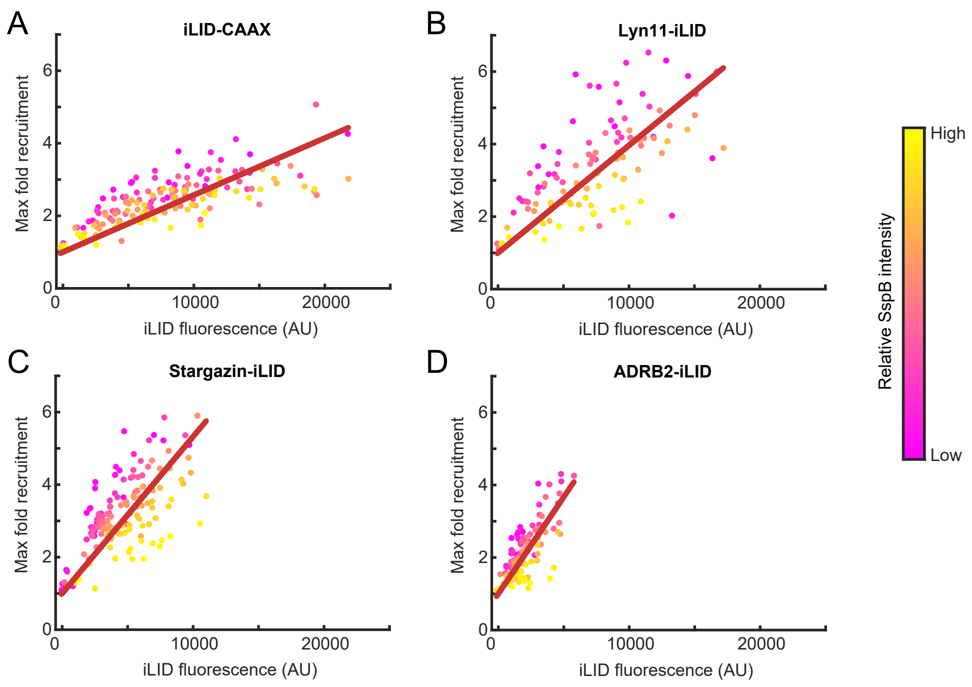

A
iLID-CAAX
B
Lyn11-iLID
C
Stargazin-iLID
D
ADRB2-iLID
Max fold recruitment
iLID fluorescence (AU)
0
2
4
6
0
10000
20000
Relative SspB intensity
High
Low

### Figure S7

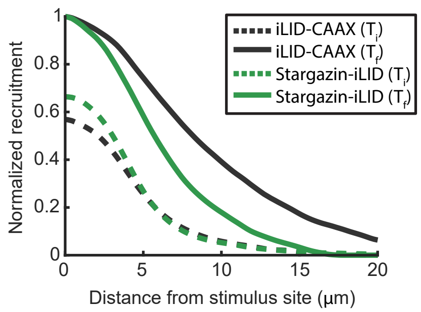

iLID-CAAX ($T_i$)
iLID-CAAX ($T_f$)
Stargazin-iLID ($T_i$)
Stargazin-iLID ($T_f$)
Normalized recruitment
0
0.2
0.4
0.6
0.8
1
0
5
10
15
20
Distance from stimulus site (μm)

### Figure S8

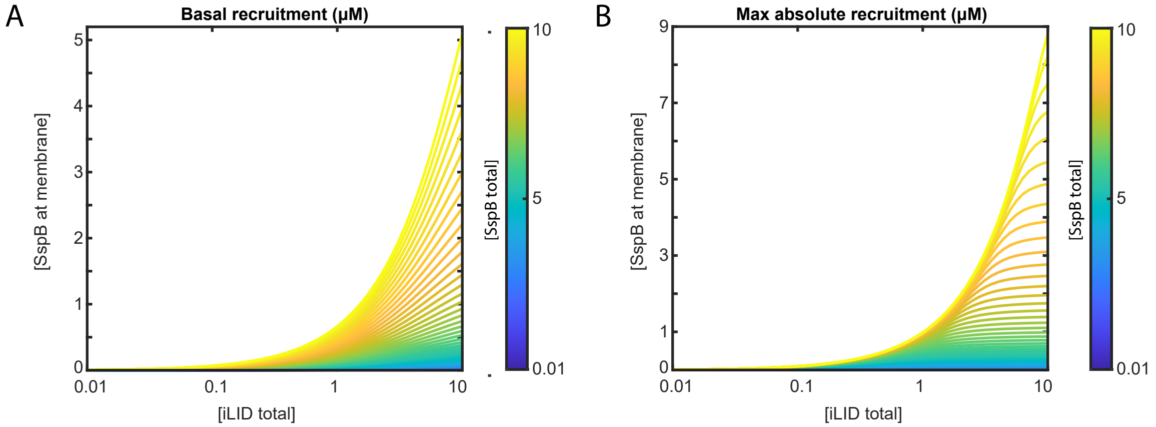
A
Basal recruitment (μM)
[SspB at membrane]
0
1
2
3
4
5
0.01
0.1
1
10
[iLID total]
[SspB total]
10
5
0.01
B
Max absolute recruitment (μM)
[SspB at membrane]
0
1
3
5
7
9
0.01
0.1
1
10
[iLID total]
[SspB total]
10
5
0.01
